# A SUMO-interacting motif in the guanine nucleotide exchange factor EPAC1 is required for subcellular targeting and function

**DOI:** 10.1101/2025.05.02.651912

**Authors:** Wenli Yang, Fang Mei, Wei Lin, Jason E. Lee, Si Nie, Christopher J. Bley, André Hoelz, Xiaodong Cheng

## Abstract

Exchange protein directly activated by cAMP (EPAC1), a multifunctional intracellular cAMP receptor, dynamically localizes to various cellular loci, engaging with diverse molecular partners to maintain cellular homeostasis. The study investigates the role of the SUMO interacting motif (SIM) in the subcellular targeting and cellular functions of EPAC1. It reveals that the SIM is a critical structural element for EPAC1’s association with RanBP2/Nup358, a nucleoporin of the cytoplasmic filament component of the nuclear pore complex (NPC). Mutational disruption of EPAC1 SIM interferes with EPAC1’s ability to activate its canonical effectors, small GTPases, Rap1 and Rap2, and non-canonical functions, such as the formation of nuclear condensates and cellular SUMOylation. Because SIM is also directly involved in cAMP binding, RanBP2’s association with EPAC1 with the SIM attenuates EPAC1’s cAMP binding affinity to generate an EPAC1 signaling microdomain with reduced cAMP sensitivity around the NPC. The coupling between EPAC1’s scaffold association and cAMP binding enables EPAC1 to tune its sensitivity to stress stimuli spatially depending on the cellular locations. These findings provide novel structural insights into EPAC1 signaling, highlighting the importance of SIM in EPAC1’s cellular functions and potential novel strategies for therapeutically targeting EPAC1.

## Introduction

Biological processes and communications are highly coordinated and tightly regulated within subcellular compartments despite the seemingly chaotic cellular milieu. It is well documented that signal transduction mediated by the diffusible small molecule second messenger cAMP occurs at localized signaling microdomains. Within these subcellular compartments, the synthesis and degradation of cAMP and interactions among cAMP receptors and their effectors are well choreographed to produce precise cellular readouts in a spatial and temporal manner (1). For example, while mammalian exchange protein directly activated by cAMP, EPAC1, is broadly expressed in different tissues (2) and has been observed at various subcellular loci in a cell-cycle dependent manner (3–7), it forms distinct signalosomes to regulate various biological processes in response to cellular stresses (8–12).

EPAC1 is a multidomain protein composed of a disordered N-terminal tail, followed by a Disheveled/Egl-10/pleckstrin (DEP) domain, a cAMP binding domain, a RAS exchange motif (REM), a RAS-association (RA) domain, and a CDC25 homology domain (CDC25-HD) (2). Cellular localization analyses have provided major insights into understanding the structure and function of EPAC1. The recognition of the importance of the DEP domain in EPAC1’s PM association function (3, 13, 14) led to the subsequent identification of a hidden 17-aa polybasic phosphatidic acid binding motif (15) and an adjacent G protein-coupled receptor kinase 2 (GRK2) phosphorylation site (S108) within the DEP domain (16). cAMP-induced conformational changes within the DEP domain (17) result in the exposure of this lipid binding motif to promote the PM recruitment of EPAC1 (15). In contrast, phosphorylation of S108 by GRK2 inhibits cAMP-induced translocation of EPAC1 to the PM, where EPAC1 signaling sensitizes the mechanosensor Piezo2 and contributes to inflammatory mechanical hyperalgesia (16). The dynamic association of EPAC1 with the PM compartment is critical for many of EPAC1’s cellular functions, such as cell adhesion (18), TSH-induced thyroid cell proliferation (19), inhibition of neurite outgrowth (20), endothelial annexin A2 cell surface translocation and plasminogen activation (21), and pathological angiogenesis through the suppression of Notch signaling (22). Likewise, the discovery of EPAC1 mitochondrial localization and the identification of an N-terminal mitochondrial-targeting sequence (3) foretells potential functional roles of EPAC1 signaling microdomains in this subcellular organelle. Indeed, ensuing studies have revealed that EPAC1 regulates a diverse array of mitochondrial functions and that their dysregulation contributes to various pathogenesis associated with mitochondrial disorders (23–26).

Despite extensive studies, significant gaps in our understanding of EPAC1 subcellular targeting and their associated functions persist. Whereas a significant fraction of cellular EPAC1 resides at the nuclear envelope (NE) and inside the nucleus (3, 6, 7, 18, 20, 27–29), the structural element/motif responsible for EPAC1’s nuclear targeting is unknown, and very little is known about EPAC1’s functions in this major cellular compartment. A study by Storck and colleagues first showed that EPAC1 interacted with Ran:GTP using the RA domain. A potential interaction between EPAC1 and RanBP2 at the NE was also proposed (6). RanBP2, also known as nucleoporin 358 (Nup358), is the largest Nup with several modular domains connected by flexible linkers or FG repeats (30, 31). It is a principal constituent of the cytoplasmic filaments of the nuclear pore complex (NPC) (32–34). A subsequent report by Bos and colleagues confirmed the interaction between EPAC1 and RanBP2 and purported that RanBP2’s Zinc finger domain (ZFD) interacted with EPAC1’s CDC25-HD domain and inhibited it activity at the NPC. This interaction was not affected by increases in intracellular cAMP as cAMP binding does not affect conformation of the CDC25-HD domain (29). However, Yarwood and colleagues later challenged this notion. They were not able to validate a direct interaction between RanBP2 and the CDC25-HD of EPAC1. In contrast, they identified a nuclear pore localization signal (amino acids 764–838) within the CDC25-HD domain of EPAC1 that appeared to be responsible for nuclear tethering by interacting with other protein constituents of the NPC (7). Therefore, the structural element in EPAC1 that is responsible for RanBP2 interaction and the mechanism of EPAC1 activation at the NPC remain unclear. In addition, the CDC25-HD/ZFD interaction model raises one key question: How does cAMP activate EPAC1 trapped by RanBP2 on the NE, one of the primary cellular EPAC1 loci?

In this study, we have identified a SUMO interacting motif (SIM), previously described as a critical structural component required for heat-shock induced EPAC1 SUMOylation (35), as the key structure element important for EPAC1’s interaction with RanBP2. EPAC1 SIM forms the middle β-strand of the tripartite β-sheet-like switchboard (SB) structure that is critical for connecting and maintaining the proper orientation between the regulatory and catalytic halves of EPAC1 to keep it in its autoinhibitory state (36). cAMP binding leads to major conformational changes centered around the SIM/SB, including a hinge motion and the translocation of the SIM/SB to become the "lid" of the cAMP binding pocket (17, 37). Our study reveals that EPAC1 SIM serves as a conformational switch, between RanBP2- and cAMP-bound states, to modulate EPAC1’s cellular targeting and activation. These results solve a key puzzle in our understanding of EPAC1 cellular targeting and function.

## Results

### RanBP2 is critical for the NE targeting but not the nuclear localization of EPAC1

Earlier studies reveal that EPAC1 is associated with RanBP2 (6, 29). To determine if interaction between EPAC1 and RanBP2 is necessary for EPAC1’s NE targeting, we used an inducible Nup358-knockout cell line (AID::Nup358 HCT116), in which an N-terminal auxin-inducible degron (AID) tag was inserted into genomic Nup358 loci. A previous study found that auxin treatment of the cell line resulted in a time-dependent and near-complete degradation of endogenous Nup358 within three hours without affecting other nucleoporins in Nup358’s vicinity (34). Indeed, we observed that auxin treatment of the cell line resulted in a time-dependent and near-complete degradation of the endogenous Nup358 protein within 3 hours (**Figure 1A**) while leaving the NPC intact, as demonstrated by staining with an anti-nuclear pore complex proteins antibody, mAb414 (**Figure 1B**). Control experiments using parental HCT116 cells showed that Auxin treatment did not significantly affect the endogenous levels of RanBP2 and EPAC1 (**Figure S1**). When we ectopically introduced a mNeonGreen (mNG)-tagged EPAC1 (EPAC1-mNG) in AID::Nup358 HCT116, a typical EPAC1 cellular distribution was observed, with EPAC1-mNG fluorescence signal enriched on the NE and within the nuclei at the basal condition. Acute depletion of RanBP2 by a three-hour auxin treatment led to diminished EPAC1 signals at NE but not inside the nucleus (**Figure 1B**). These results suggest that RanBP2/Nup358 is critical for the NE targeting of EPAC1.

**Figure 1.**
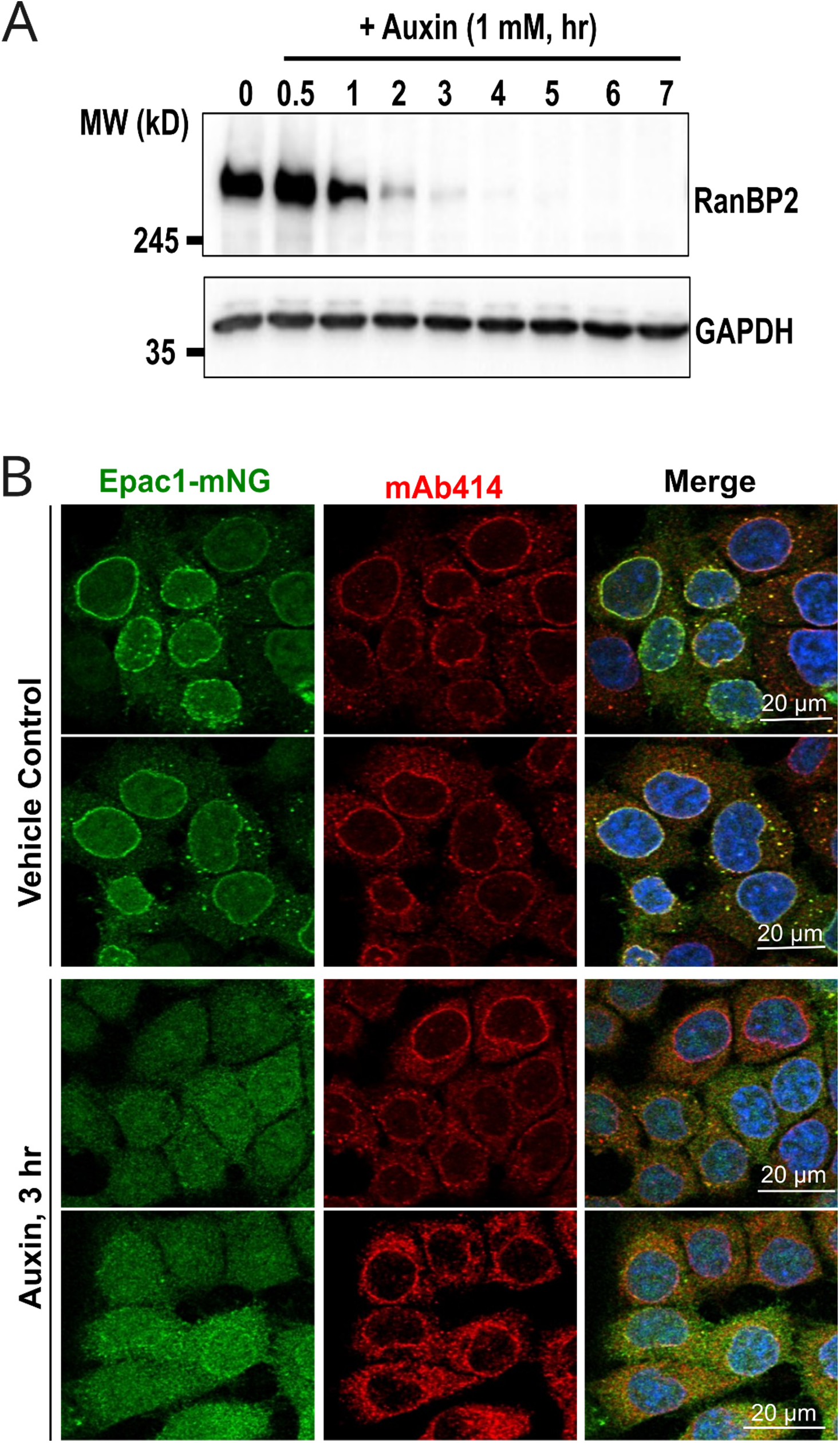
RanBP2/Nup358 is essential for NE localization of EPAC1. (A) Time-dependent depletion of endogenous RanBP2/UuP358 by auxin. The expression levels of RanBP2 probed by immunoblotting analysis using anti-RanBP2 antibody in AID::NUP358 HCT116 cells treated with 1 mM auxin at 37 °C for various times. (B) Confocal imaging of AID::Nup358 HCT116 cells ectopically expressing EPAC1-mNG and treated with vehicle or Auxin for 3 hours. mAb414 staining marks the nuclear envelope. Bar = 20 μm.

To identify the structural element/domains in RanBP2 responsible for EPAC1’s NE targeting, we ectopically expressed N-terminally 3×HA-tagged full-length (FL) RanBP2 or various C-terminal deletion variants in AID::Nup358 HCT116 cells stably expressing EPAC1- mNG (**Figure 2A**). All these RanBP2 variants can localize to the NE correctly (29). When endogenous RanBP2 was depleted by 3 hr of auxin treatment, FL and NTD-ZFD RanBP2 were able to restore the NE-targeting of EPAC1 while NTD-RanBD1 and NTD-OE RanBP2 failed to rescue (**Figure 2B**), suggesting that RanBP2 ZFD is responsible for the NE targeting of EPAC1. To determine if RanBP2 ZFD binds directly with EPAC1, we recombinantly expressed and purified the ZFD of RanBP2. Glutathione-beads pull-down of purified GST-EPAC1 incubated with RanBP2-ZFD led to robust co-precipitation of RanBP2-ZFD (**Figure 2C**). These results suggest that the ZFD of RanBP2 binds directly with EPAC1 and is necessary for its NE targeting.

**Figure 2.**
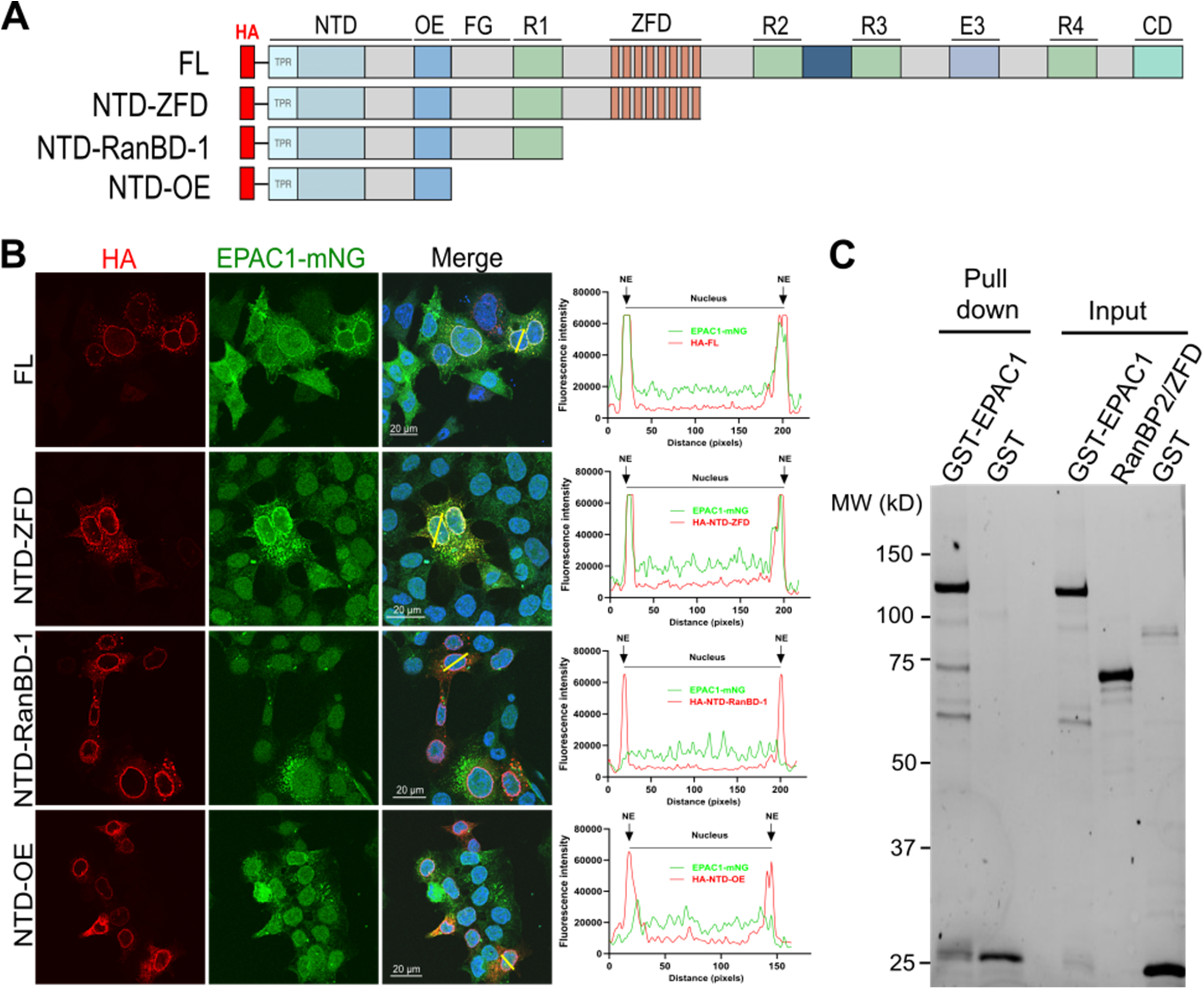
The Zinc Finger Domain of RanBP2/Nup358 is responsible for the NE localization of EPAC1. (A) Domain structures of the transfected Nup358 variants. (B) Subcellular localization of N-terminally 3×HA-tagged Nup358 variants (Red) including Nup358 full length (FL), NTD-ZFD, NTD-RanBD-1, NTD-OE and EPAC1-mNG in AID::Nup358 HCT116 cells visualized by immunofluorescence confocal imaging after auxin-induced deletion of endogenous RanBP2. Bar = 20 μm. Line-scan plots show the quantified fluorescence intensities of EPAC1-mNG (green) and HA-tagged Nup358 variants (red) along the yellow lines crossing the nucleus, analyzed using ImageJ software. (C) Protein gel showing purified GST- Epac1 pulled down with RanBP2-ZFD using glutathione beads.

### A SUMO interacting motif (SIM) of EPAC1 is essential for RanBP2 interaction

To identify the structural element in EPAC1 important for RanBP2 interaction, we performed affinity pulldown analyses using GFP-tagged full-length wild-type EPAC1, a cAMP-binding deficient mutant, EPAC1(R279E), N-terminal regulatory and C-terminal catalytic halves of EPAC1 (EPAC1(1–330) and EPAC1(331–881)). To our surprise, while both the full-length EPAC1 and R279E mutant were able to pull down RanBP2 robustly with the cAMP binding deficient mutant pulling down slightly more RanBP2, neither EPAC1(1–330) nor EPAC1(331–881) interacted with RanBP2 (**Figure 3B**). These results suggest either that both the regulatory and the catalytic halves of EPAC1 are required for RanBP2 association or that a properly oriented regulatory and the catalytic halves of EPAC1, i.e., the linker, is essential for EPAC1 and RanBP2 interaction. Indeed, when we mutated the recently identified SIM motif (35), five key residues ^320^VVLVL^324^ connecting the regulatory and the catalytic regions to alanines, the EPAC1(SIM5A) mutant was not able to interact with RanBP2 (**Figure 3B**). We further validated our finding by performing reverse affinity pulldown analysis. Significantly less EPAC1(SIM5A) was observed to co-precipitate with RanBP2 (**Figure 3C**). Moreover, we generated more conservative EPAC1 SIM mutants, EPAC1(V323A/L324A) or EPAC1(V321A/V323A). Affinity pulldown analyses showed that the ability of EPAC1(SIM2A) mutants to interact with RanBP2 was dramatically reduced (**Figure 3D**). These data suggest that EPAC1-SIM is an important structural element for EPAC1 and RanBP2 interaction.

**Figure 3.**
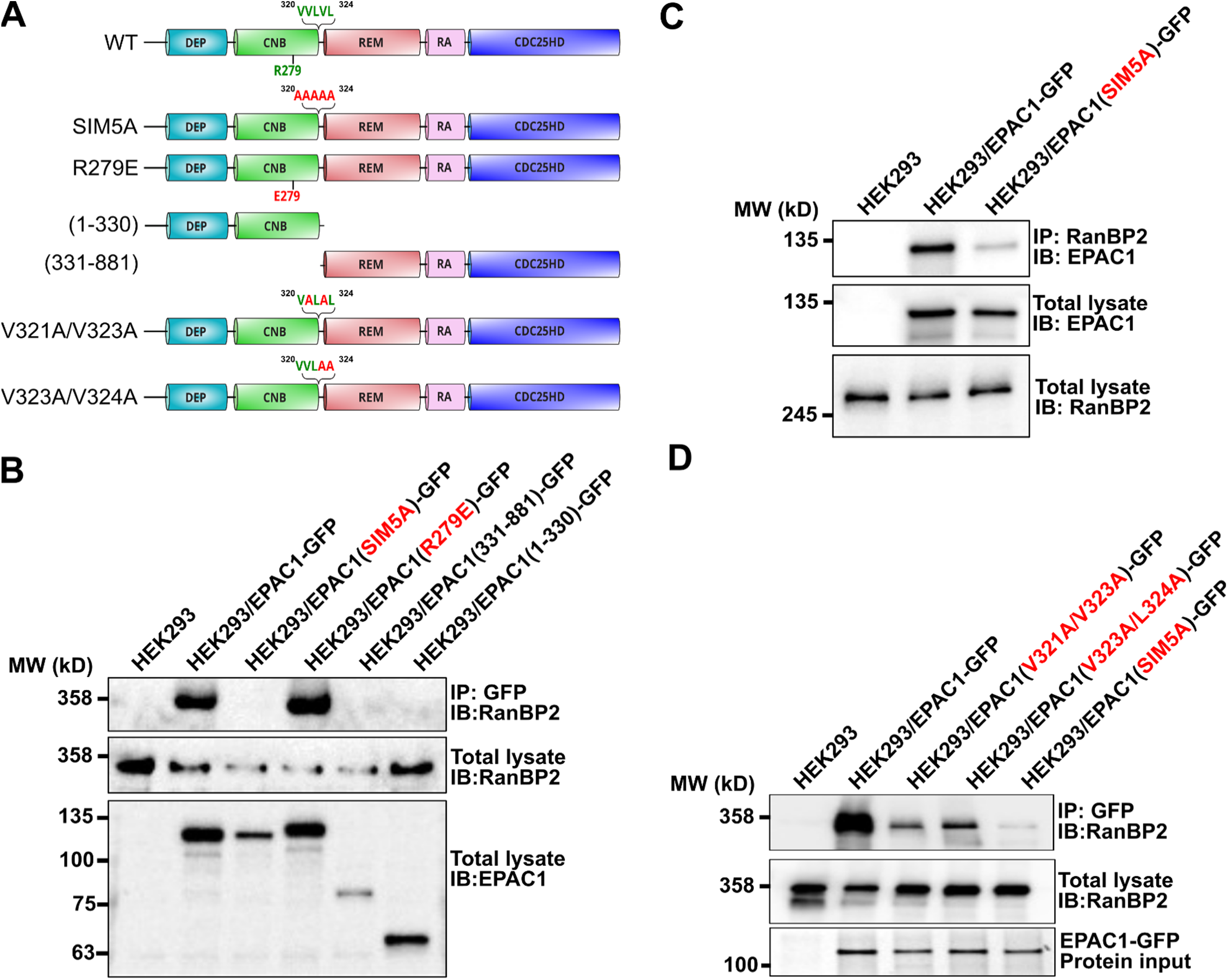
EPAC1 SIM element is critical for EPAC1 and RanBP2 cellular interaction. (A) The Domain structure of transfected EPAC1 variants. (B) Association of endogenous RanBP2 with ectopically expressed EPAC1 constructs tagged with a C-terminal GFP in HEK293 cells probed by immunoprecipitation using an anti-GFP antibody. (C) Association of ectopically expressed GFP-tagged EPAC1 constructs with endogenous RanBP2 in HEK293 cells probed by immunoprecipitation using an anti-RanBP2 antibody. (D) Association of endogenous RanBP2 with ectopically expressed EPAC1 SIM variants tagged with a C-terminal GFP in HEK293 cells probed by immunoprecipitation using anti-GFP antibody.

### RanBP2 ZFD competes with cAMP for SIM binding and inhibits EPAC1 GEF activity

Considering the important role of SIM/SB plays in EPAC1 activation, we hypothesize that binding of ZFD may suppresses cAMP-induced EPAC1 GEF activity and that cAMP-induced change in SIM conformational state may disrupt the interaction between EPAC1 and RanBP2 in cells. Indeed, when incubated with EPAC1, RanBP2-ZFD inhibited cAMP induced EPAC1’s GEF activity while having minimal effects on EPAC2’s GEF activity (**Figure 4A)**. Furthermore, when titrated with various cAMP concentrations, RanBP2-ZFD shifted the AC_50_ value of cAMP to higher concentration by roughly 10 folds and reduced the maximal GEF activity significantly (**Figure 4B)**.

**Figure 4.**
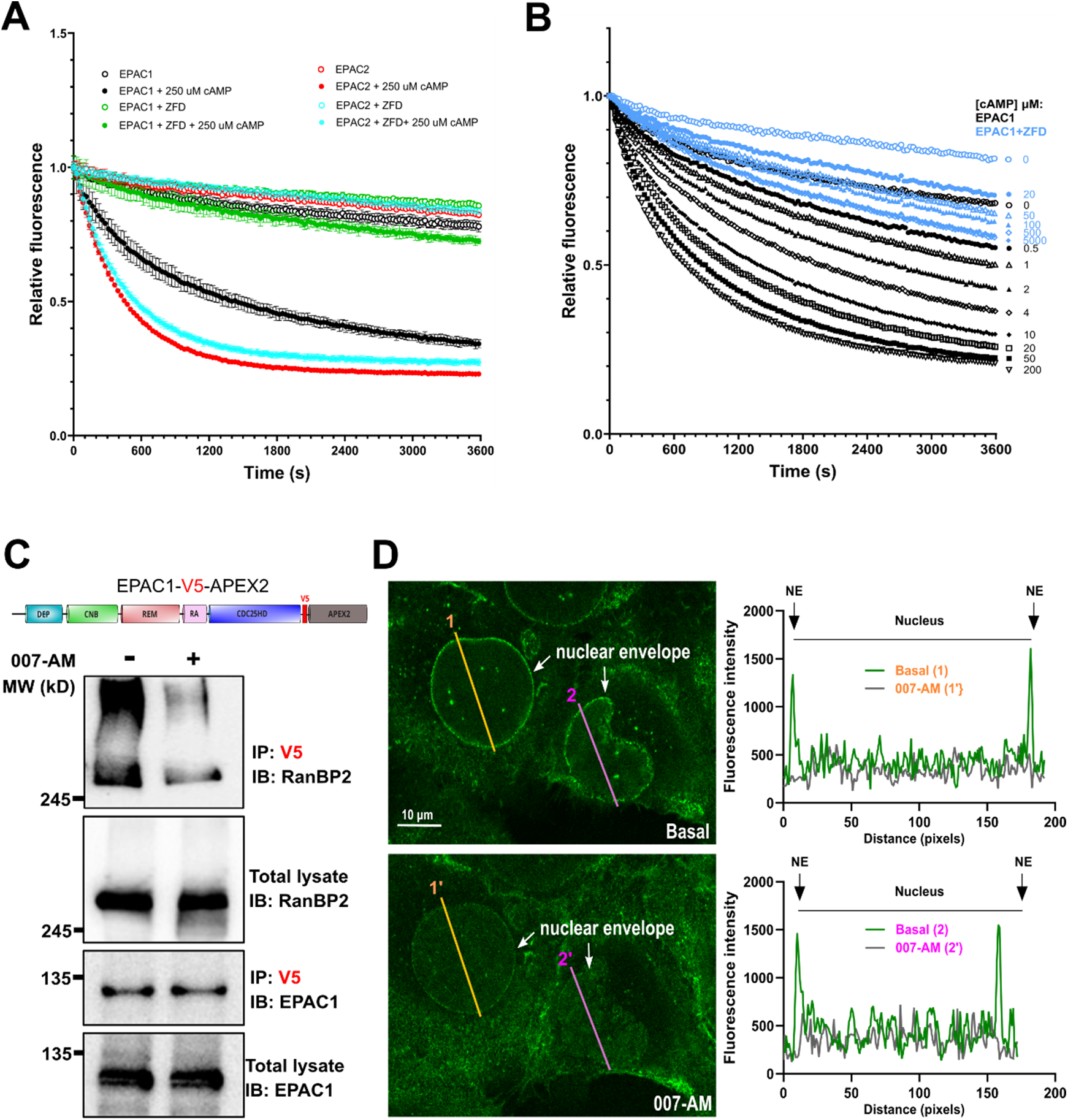
RanBP2 ZFD inhibits EPAC1 activation and EPAC1 activation by cAMP reduces its cellular interaction with RanBP2. (A) In vitro basal and cAMP induced EPAC1 and EPAC2 GEF activity monitored in the presence or absence of RanBP2 ZFD. (B) Dose-dependent activation of EPAC1 by cAMP in the absence or presence of RanBP2 ZFD. (C) cAMP binding reduces EPAC1 and RanBP2 cellular interaction. Association of endogenous RanBP2 with EPAC1 in HEK293 cells stably expressing EPAC1-V5-APEX2 probed by immunoprecipitation using anti-V5 magnetic beads in response to 007-AM treatment. (D) 007-AM (5 μM) treatment reduces EPAC1 nuclear envelope localization. Snapshots and line-scan plots of live cell fluorescence confocal imaging of EPAC1-mNG in U2OS cells under basal and 007-AM treatment conditions. Bar = 10 μm.

To test if cAMP competes with RanBP2 for binding EPAC1 in cells, we performed affinity purification of ectopically expressed EPAC1-V5-APEX2 in HEK293 cells using anti-V5 magnetic beads. As shown in **Figure 4C**, 007-AM treatment led to a significant reduction of co-purified endogenous RanBP2. Moreover, when we ectopically expressed EPAC1-mNG in U2OS cells and monitored the effect of EPAC1 activation on its cellular localization using confocal live cell imaging, EPAC1-mNG was observed to be enriched at NE as expected. The administration of an EPAC1-specific agonist, 007-AM, led to a significant reduction in the NE association of EPAC1- mNG with concomitant redistribution of EPAC1-mNG in the cytoplasm and nucleus (**Figure 4D**). In sum, these results suggest that RanBP2-ZFD interacts with EPAC1 in its autoinhibitory conformation and that cAMP binding disrupts the interaction between EPAC1 SIM and RanBP2 ZFD.

### Mutation of EPAC1 SIM suppresses cAMP-induced EPAC1’s cellular guanine nucleotide exchange activity

Proper subcellular compartmentalization of EPAC1 is critical for its cellular functions. Considering the importance of EPAC1 SIM in interacting with RanBP2 at the NE, a major cellular locus of EPAC1, we hypothesized that EPAC1 SIM might play a role in EPAC1’s canonical cellular guanine nucleotide exchange activity. To test this hypothesis, we compared the ability of EPAC1(SIM5A) or WT EPAC1 to activate cellular Rap1 and Rap2 in response to cAMP by monitoring the cellular Rap-GTP levels using a glutathione S-transferase fusion of the Rap1-binding domain of RalGDS that preferentially binds with GTP-bound Rap GTPases. As shown in **Figure 5**, activation of cellular EPAC1 by 007-AM led to robust increases in cellular Rap1 and Rap2 in HEK293 cells, stably expressing EPAC1-APEX2. On the other hand, the levels of Rap1-GTP and Rap2-GTP were mostly unaltered in HEK293 cells, stably expressing EPAC1(SIM5A)- APEX2. Similar results were observed when cells were treated with isoproterenol (ISO), a selective agonist of the β-adrenergic receptors (**Figure S2**). These results suggest that mutation of EPAC1 SIM disrupts its ability to mediate cAMP’s effect of activating Rap GTPases.

**Figure 5.**
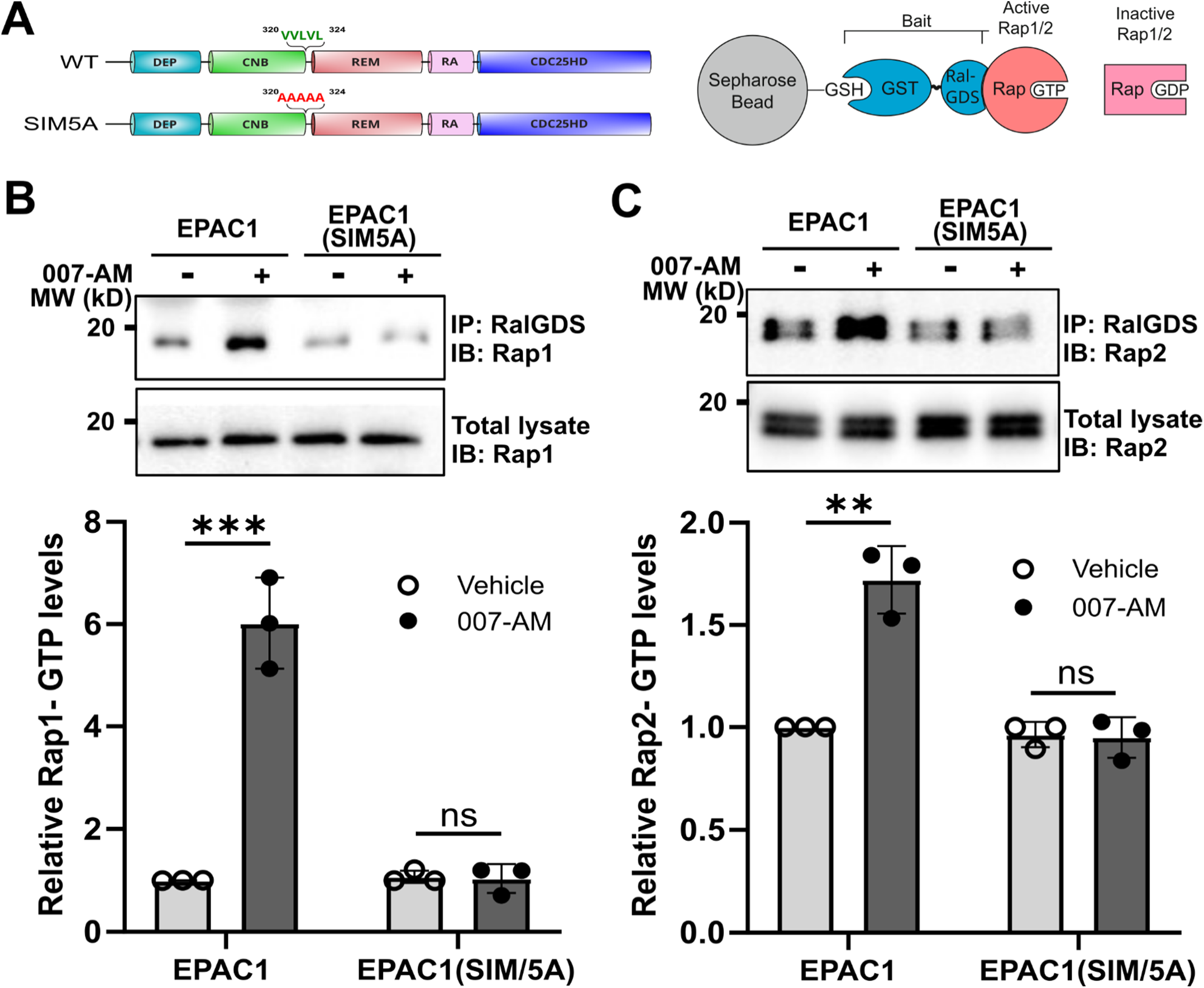
EPAC1 SIM mutation interferes with cAMP-induced cellular activation of Rap small GTPases. (A) Schematic representation of the GST-RalGDS mediated active Rap GTPase pulldown. Levels of cellular Rap1-GTP (B) or Rap2-GTP (C) in HEK293 cells ectopically expressing EPAC1-EYFP or EPAC1(SIM5A)-EYFP in response to 007-AM treatment. Data are presented as Mean ± SEM (N = 3).

### EPAC1 SIM is required for cAMP-induced EPAC1 nuclear condensates and cellular SUMOylation

Recent studies have revealed that EPAC1 activation by cAMP promotes the formation of nuclear condensates enriched with EPAC1 and components of the SUMOylation machinery and, consequently, leads to an enhanced cellular SUMOylation in a Rap-independent manner (38). Furthermore, we have also demonstrated that EPAC1 can undergo SUMOylation itself at K561 in response to heat shock and that SIM plays critical roles in interacting with SUMO-conjugating enzyme UBC9 and heat shock-mediated EPAC1 SUMOylation and nuclear condensate formation (35). These findings prompt us to ask if EPAC1 SIM also plays a role in forming the nuclear condensates and regulating cellular SUMOylation in response to cAMP. When HEK293 cells ectopically expressing EPAC1-YFP or EPAC1(SIM5A)-YFP were subjected to 7 min stimulation with 5 μM 007-AM or 20 μM ISO, robust formation of nuclear condensates was observed (**Figure 6 A & B**). On the other hand, EPAC1(SIM5A)-YFP could not form nuclear condensates in response to 007-AM or ISO treatment (**Figure 6 C & D**). In addition, it appears that the overall levels of nuclear EPAC1(SIM5A)-YFP were significantly reduced as compared to EPAC1-YFP, suggesting that SIM of EPAC1 plays a role in the nuclear targeting of EPAC1

**Figure 6.**
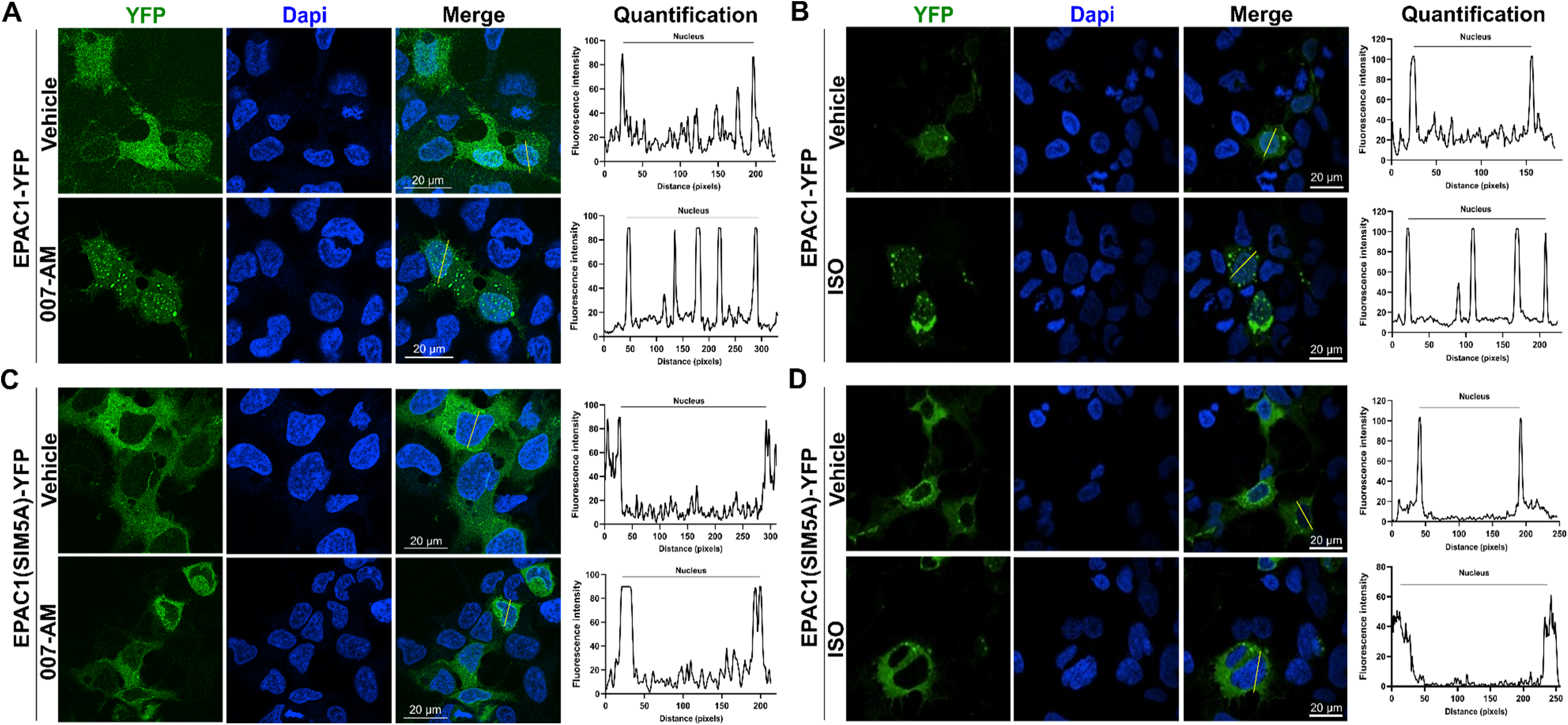
EPAC1 SIM element plays an essential role in cAMP-induced EPAC1 nuclear condensate. Confocal images of HEK293 cells ectopically expressing EPAC1-EYFP in response to 5 μM 007-AM (A) or 20 μM ISO (B). Confocal images of HEK293 cells ectopically expressing EPAC1(SIM5A)-EYFP in response to 007-AM (C) or ISO (D). Bar = 20 μm. Line-scan plots show the quantified fluorescence intensities of EPAC1-YFP along the yellow lines crossing the nucleus, analyzed using ImageJ software.

Consistent with these findings associated with nuclear condensate formation in response to heat shock and 007-AM, when HEK293 cells stably expressing EPAC1-APEX2 and EPAC1(SIM5A)-APEX2 were subjected to the same treatments, we observed robust increases in cellular SUMOylation in EPAC1-APEX2 cells under both heat shock and 007-AM conditions, while levels of cellular SUMOylation in EPAC1(SIM5A)-APEX2 cells remained unaffected (**Figure 7 A&B**). After the heat shock treatment at 43 °C for 30 min, the basal EPAC1 levels decreased slightly due to EPAC1 SUMOylation (35) and recovered after returning to 37 °C for 2 hr (**Figure S3**). Taking together, these observations indicate that EPAC1 SIM is a key structural motif for EPAC1’s functions involving nuclear condensate formation and regulation of cellular SUMOylation.

**Figure 7.**
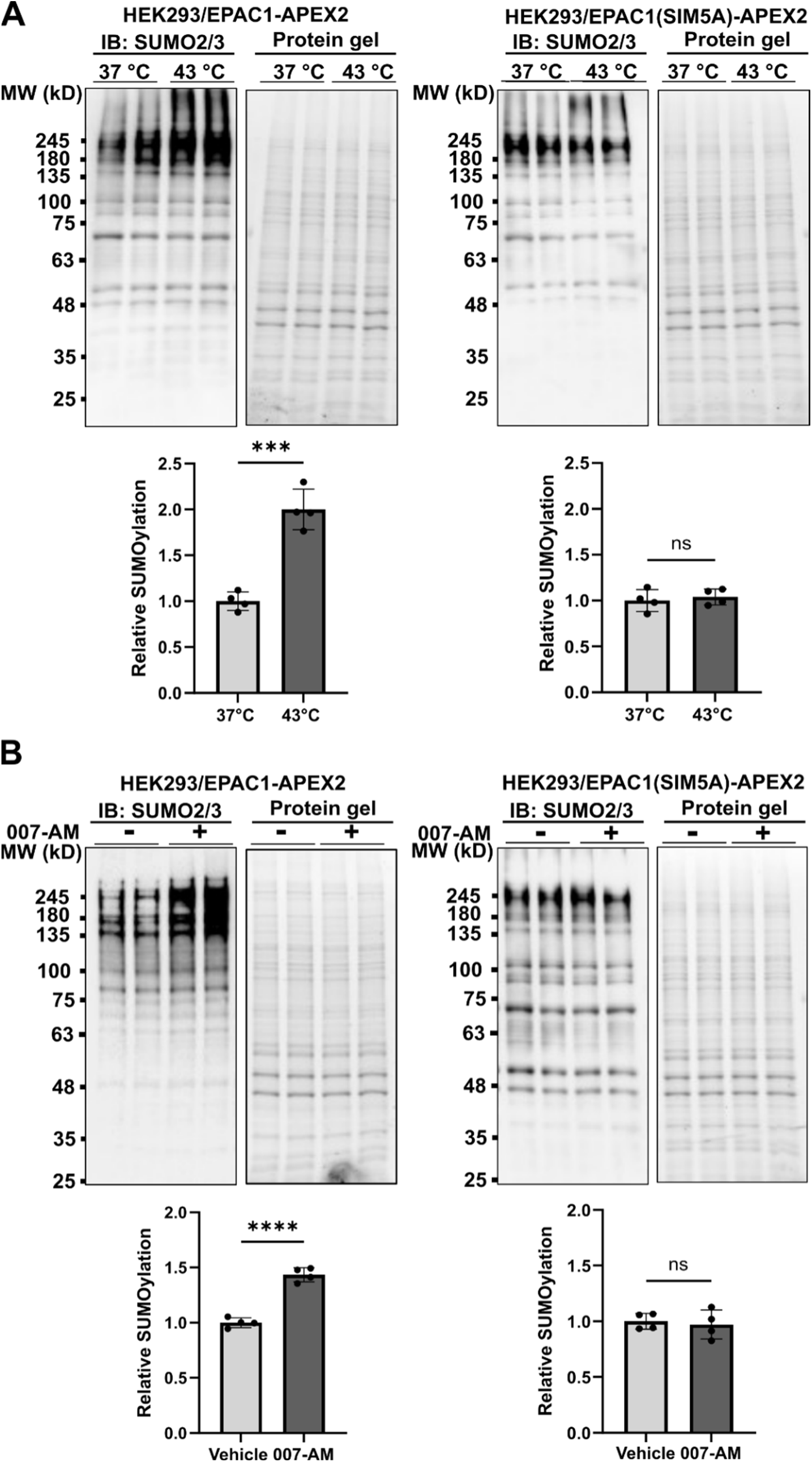
EPAC1 SIM mutation interferes with cAMP and heat shock-induced cellular SUMOylation. Immunoblotting showing cellular SUMOylation levels in HEK293 cells stably expressing EPAC1-APEX2 and EPAC1(SIM5A)-APEX2, detected by anti-SUMO2 antibody in response to heat shock treatment (A) or 5 μM 007-AM (B) treatment. Data are presented as Mean ± SEM (N = 4).

## Discussion

Subcellular localization analyses of EPAC1 have provided valuable insights into our understanding of its versatile cellular and physiological functions. Although EPAC1 has been observed to be enriched on the NE and reside within the nucleus for over two decades, our understanding of its targeting mechanism and functions in this important subcellular compartment is scarce. In this study, we identified a SIM as a key structural element for EPAC1’s interaction with RanBP2 and nuclear targeting. This finding fills in a significant gap in the current mechanistic understanding of EPAC1’s subcellular targeting and functions. Previous work proposed that RanBP2 interacted with EPAC1’s CDC25-HD catalytic domain and inhibited its activity at the nuclear pore, and this interaction was in sensitive to cAMP because cAMP binding does not change the conformation of the CDC25-HD domain (29). A key problem with this model is that EPAC1 trapped by RanBP2 at the NE, one of the primary cellular EPAC1 loci, will not be able activated by cAMP. In contrast, a SIM-based RanBP2 interaction is tunable by cAMP and structurally reasonable. In our model, RanBP2 interacts with SIM of EPAC1 in its inactive conformation as cAMP-binding deficient EPAC1 mutant binds RanBP2 better than WT EPAC1 (**Figure 3B**). When intracellular cAMP increases, cAMP binding leads to conformational changes of the SIM, which disrupts interaction with RanBP2 and leads to EPAC1 activation and its dissociation from the NE (**Figure 4**).

The involvement of SIM in both cAMP and RanBP2 binding has broad implications for EPAC1 cellular functions, particularly for its compartmentalized signaling. Inactive EPAC1 tethered at NE by RanBP2 will have an increased activation threshold than free EPAC1 in the cytosol, as cAMP must compete with RanBP2 for EPAC1 binding (**Figure 4**). This is opposite to a PM-associated EPAC1, where a recent study shows that PM primes EPAC1 for cAMP binding and dramatically decreases its activation threshold (39). Therefore, depending on its cellular localization and interacting partners, cellular EPAC1 proteins will have different sensitivities for cAMP activation with a spatial gradient of high, medium, and low, starting from the PM and ending at the NE/NPC. As such, the dynamic range of cellular EPAC1 signaling is significantly expanded as more intense stimuli will be required to propagate the EPAC1 signaling from the cell surface to the nucleus (**Figure 8**). Moreover, considering the dynamic nature of EPAC1 subcellular localization, such a mechanism also endows cells to rewire the sensitivity of the cellular EPAC1 signaling network via redistribution of cellular EPAC1 in response to environmental stimuli.

**Figure 8.**
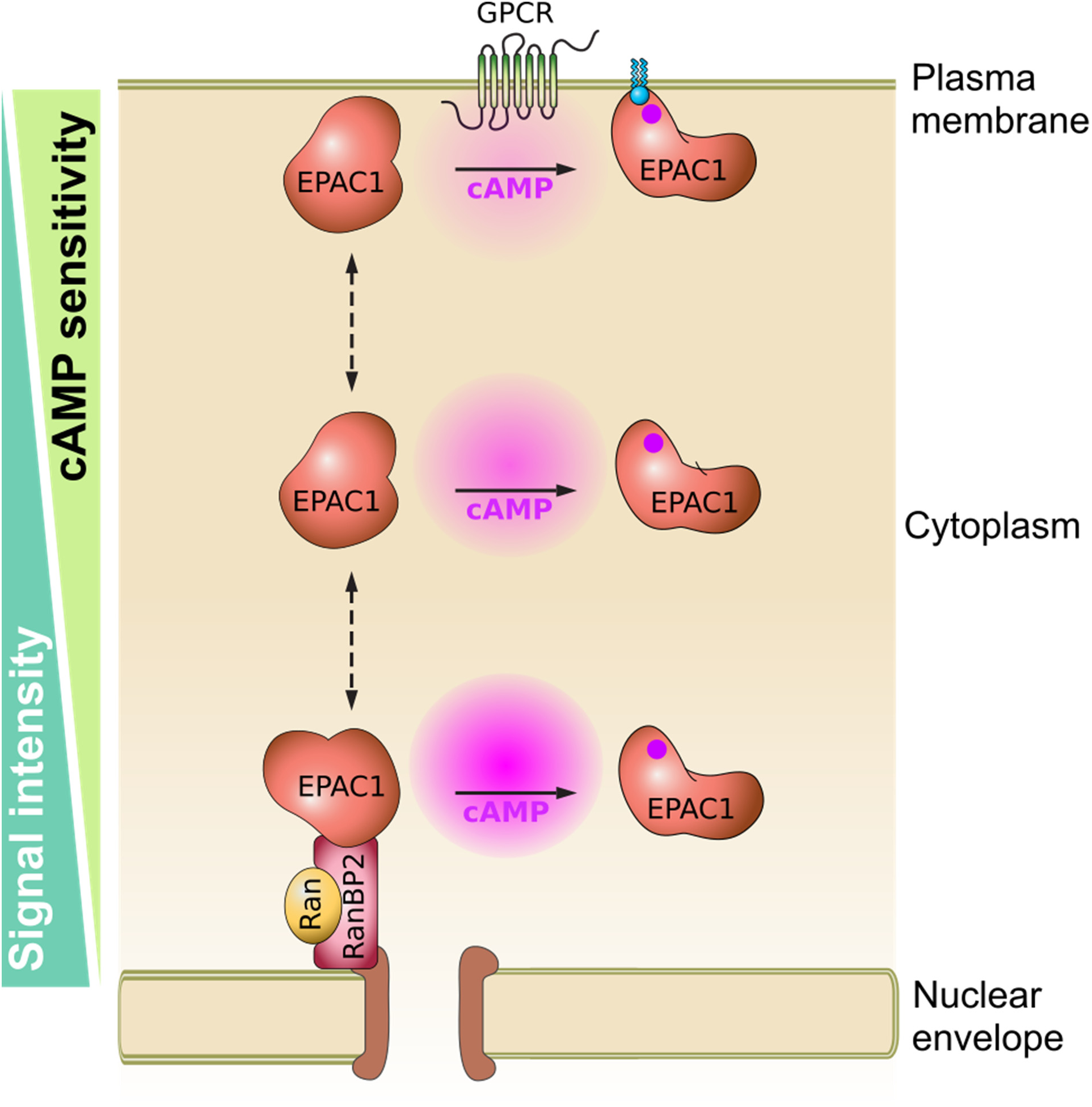
Schematic model of intracellular EPAC1 signaling sensitivity gradient. Interaction of EPAC1 with lipids on the plasma membrane or RanBP2 associated with the nuclear pore complex prime or impedes cAMP binding to EPAC1 at the respective cellular locus.

Proper subcellular localization of EPAC1 is essential for its cellular activation and functions. Indeed, the ability of the EPAC1 SIM mutants to activate its downstream canonical effectors, small GTPases Rap1 and Rap2, in response to cAMP is significantly diminished. In addition, EPAC1 SIM mutants do not form nuclear condensates and are compromised in promoting cellular SUMOylation in response to both heat shock and ligand stimulation. Our previous studies have shown that EPAC1-mediated cellular SUMOylation does not involve Rap1 and 2 but involves the formation of nuclear condensates. Therefore, these observations suggest that EPAC1 SIM is important for both canonical and nonconical EPAC1 functions. This functional duality of the SIM in ligand-induced activation and SUMOylation regulation has important implications. At the molecular level, the SIM region is likely co-evolved for cAMP binding and SUMO recognition and acts as a key structural element linking these two major stress response pathways. Under this notion, when examining EPAC1 biological functions, we need to keep in mind that cAMP signaling and SUMOylation are likely cross-modulating each other, as we have shown in our recent studies (35, 38).

On the translational side, SIM provides an ideal target for designing pharmacological agents capable of modulating both canonical and nonconical EPAC1 functions. This is important because while several conventional EPAC-specific small molecule pharmacological probes are available for modulating canonical EPAC1 functions, they are likely not as effective for cellular functions associated with EPAC1 biomolecular condensates, which have been recently shown to play important roles in EPAC1 cellular functions, particularly in the nucleus. Currently, developing pharmacological agents capable of modulating biomolecular condensates remain challenging. The unique structure properties of the SIM provide a niche for developing chemical or biological agents that are capable of modulating EPAC1-specific condensates.

## Materials and Methods

### Materials and reagents

Auxin (3-Indoleacetic acid, Catalog no. I3750-5G-A), cOmplete™, EDTA-free Protease Inhibitor Cocktail (Catalog no. 4693132001), isoproterenol hydrochloride (Catalog no. I5627), Dulbecco’s modified Eagle’s medium (DMEM) high glucose (Catalog no. D5796), fetal bovine serum (FBS) (Catalog no. F2442), N-ethylmaleimide (NEM) (Catalog no. E3876), poly-llysine solution (0.01%, catalog no. P4707) and FluorSave Reagent (Catalog no. 345789) were from MilliporeSigma. Antibiotic–antimycotic (100 ×) (Catalog no. 15240096), 4′,6-diamidino-2-phenylindole (DAPI) (Catalog no. 62248), Lipofectamine 2000 (Catalog no. 11668-019) and Goat anti-Rat IgG (H+L) Cross-Adsorbed Secondary Antibody, Alexa Fluor™ 594 (Catalog no. A-11007) were from Thermo Fisher Scientific. Protein A/G PLUS-Agarose (Catalog no. sc-2003) was from Santa Cruz Biotechnology Inc. V5-Trap® Agarose (Catalog no. v5ta) were from Proteintech Group, Inc. 8- (4-Chlorophenylthio)-2′-O-methyladenosine-3′,5′-cyclic monophosphate acetoxymethyl ester (007-AM) was from Axxora (Catalog no. BLG-C051).

### Constructions

pcDNA3.1 vectors with N-terminally 3×HA-tagged full-length RanBP2 or various C-terminal deletion variants were described previously (34). RanBP2 ZFD domain (1171-1809 aa) was subcloned into the pGEX-4T-2 vector between the BamH1 and Not1 sites in frame with the N-terminal GST tag. Wild-type and mutant Human EPAC1 tagged with a C-terminal EYFP was described previously (3). Wild-type and mutant EPAC1 fused with a V5 and an APEX2 tags at the C-terminal were cloned into a lentivector, pCDH-CMV-MCS-EF1α-Puro, as described previously (35). PCR amplified EPAC1b was digested with HindIII and EcoRI and inserted into an LR Clonase^TM^ competent entry vector with the multiple cloning sites upstream of *mNG*. *EPAC1b-mNG* was then transferred to a lentiviral destination plasmid with an LR Clonase reaction. The lentiviral destination plasmid (pHAGE-pgk-cFLAG-HA-dest_puro) is driven by a mouse phosphoglycerate kinase (*mPGK1*).

### Cell culture, transfection, and stable cell lines

HEK-293 were grown in DMEM supplemented with 10% fetal bovine serum (FBS), U2OS, and AID::Nup358 HCT116 cells (34) were grown in McCoy’s 5A medium containing 10% fetal bovine serum (FBS), 1× antibiotic–antimycotic at 37°C in a 5% CO_2_ humidified incubator. All cell transfections were performed using Lipofectamine 2000 according to the manufacturer’s instructions. AID::Nup358 HCT116 and U2OS cells stably expressing EPAC1-mNG or HEK-293 cells stably expressing EPAC1-V5-APEX2 or EPAC1(SIM5A)-V5-APEX2 were established as described previously (35). Briefly, HEK293T cells at 70% confluence were transfected with pHAGE-EPAC1-mNG, pCDH-EPAC1-V5-APEX2 or pCDH-EPAC1(SIM5A)-V5-APEX2 plus the MISSION Lentiviral Packaging Mix (Sigma-Aldrich, catalog no. SHP001 and SHP002) using Lipofectamine 2000. Viral supernatant was harvested at 48 hours post-transfection and utilized to infect AID::Nup358 HCT116, U2OS, or HEK293. Stable cell lines expressing moderate EPAC1-mNG or EPAC1–V5-APEX2 were selected and used for subsequent experiments.

### Recombinant RanBP2 ZFD expression and purification

pGEX-4T-2 vector with RanBD-ZFD insert was transformed into *E. coli* strain CK600K and grown in LB supplemented with 50 µg/ml Kanamycin, 100 µg/ml ampicillin and 100 µM ZnSO_4_. Protein expression was induced with 0.1 mM IPTG overnight at 21 °C. GST-fusion recombinant proteins were purified using Glutathione Sepharose 4B resin. The GST-tag was removed by thrombin cleavage and RanBP2 ZFD was further purified by a FPLC Superdex 200 Increase gel filtration column equilibrated with 50 mM Tris, pH7.5, 150 mM NaCl, 10 µM ZnCl_2_, 1 mM EGTA, and 1 mM TCEP.

### Interaction between EPAC1 and RanBP2 ZFD accessed by affinity pulldown

Purified recombinant GST-EPAC1 protein (>90% purity, 0.3 nMol) or GST alone was immobilized on glutathione-agarose beads (50 µL of a 50% slurry per sample) at 4°C in buffer A (50 mM HEPES, pH 7.1, 250 mM NaCl, 1 mM MgCl₂, 20 µM ZnCl₂, 0.5% C12E9, 1 mM TCEP, and 1× protease inhibitor cocktail). Following immobilization, an equal amount (0.3 nMol) of RanBP2-ZFD was added to each sample, and the mixtures were gently rotated for 45 minutes at 4 °C. Beads were then washed twice with buffer A and three times with buffer B (identical to buffer A but containing 0.05% C12E9 instead of 0.5%). Bound proteins were eluted using 20 mM reduced glutathione (GSH) in 45 µL of buffer B, followed by the addition of 15 µL of 4× Laemmli sample buffer. Samples were then analyzed by SDS-PAGE to assess protein interactions

### EPAC1 subcellular localization in response to RanBP2/Nup358 depletion

AID::Nup358 HCT116 cells stably expressing EPAC1-mNG plated on glass coverslips coated with Poly-L-lysine (10 μg/ml) in 12-well plates overnight were treated with 1 mM auxin or vehicle. Following 3 hours of incubation, the media was removed, and cells were washed in PBS and then fixed in PBS supplemented with 4% paraformaldehyde (PFA) for 15 minutes at room temperature. After two washes with PBS, the cells were permeabilized with 0.25% Triton X-100 for 10 min, rinsed with PBS three times, incubated with 5% normal goat serum in PBS for 30 min to block nonspecific binding, followed by incubation of anti-Nuclear Pore Complex Proteins Antibody (mAb414, 1:1000 dilution, BioLegend, Catalog No. 902907) at 4 °C overnight. After washing with 1× TBST (20 mM Tris, 137 mM NaCl, 0.1% Tween 20) three times, cell specimens were incubated with CoraLite594 – conjugated Goat Anti-Mouse IgG(H+L) (1:300 dilution, Ptoteintech, Catalog No. SA00013-3) for 30 min at room temperature in the dark. Cell nuclei were stained with DAPI solution for 10 minutes after washing with 1× TBST three times. Coverslips were mounted with FluorSave reagent for fluorescence microscopic imaging with the Nikon AXR confocal microscope system using the same parameter settings. For rescuing analyses using RanBP2 variants, AID::Nup358 HCT116/EPAC1-mNG cells seeded on glass coverslips coated with Poly-L-lysine at 50% confluence were transfected with pcDNA3.1-3×HA-Nup358, pcDNA3.1-3×HA-Nup358-NTD-ZFD, pcDNA3.1-3×HA-Nup358-NTD-RanBD1 or pcDNA3.1-3×HA-Nup358-NTD-OE plasmid for 24 hours and then treated with 1 mM auxin or vehicle for 3 hours. The cells were then washed in PBS, fixed with 4% PFA as described above and permeabilized with 0.25% Triton X-100 for 10 min. After rinsing with PBS three times, the cells were incubated with 5% normal goat serum in PBS for 30 min to block nonspecific binding, followed by incubation of anti-HA High-Affinity antibody (1:200 dilution, Roche, Catalog no. 11867423001 or MilliporeSigma, Catalog no. I3750-5G-A) at 4 °C overnight. After washing with 1× TBST three times, cell specimens were incubated with Goat anti-Rat IgG (H+L) Cross-Adsorbed Secondary Antibody, Alexa Fluor™ 594 (1:200 dilution) for 30 min at room temperature in the dark. After washing with 1× TBST three times, cell nuclei were stained with DAPI solution. Coverslips were mounted with FluorSave reagent for fluorescence microscopic imaging with the Nikon AXR Confocal Microscope System. Co-occurrence of fluorescence signals was assessed using direct line analysis in ImageJ. Channels from each image were first separated using the “Split Channels” function. A straight line was then drawn through the nucleus of each cell in one channel, and the same line was replicated on the second channel using the “Restore Selection” function. Plot profiles were generated for each channel, capturing the relative fluorescence intensities along the line, and were then used to produce line graphs.

### EPAC1 nuclear condensates analysis

HEK293 were grown on Poly-L-lysine coated coverslips until ∼50% confluency and then transfected with pEYFP-EPAC1 or pEYFP-EPAC1(SIM5A) for 24 hours. For ISO treatment, cells were incubated with 20 uM ISO in complete medium for 30 minutes. For 007-AM treatment, cells were first starved in serum-free (SF) DMEM for 1 hour and then were treated with 007-AM (5μM) or vehicle control for 7 min. Cells were subsequently fixed in 4% PFA for 15 minutes and subjected to similar wash steps as described above. Coverslips were mounted with FluorSave reagent for fluorescence microscopic imaging with the Nikon AXR Confocal Microscope System. The fluorescence signals was assessed using direct line analysis in ImageJ as previously described (38).

### Confocal imaging analysis

Fluorescence microscopic images were collected using a Nikon AXR laser confocal microscope system. For individual control and treatment groups, the same system settings, including the laser power and exposure time, were applied. Images for more than eight randomly selected fields from at least three independent coverslips per treatment condition were collected. For live-cell imaging, U2OS/EPAC1-mNG cells were seeded overnight in glass-bottom plates (MatTek, catalog no. P35G-1.5-14-C) and imaged with a Nikon A1R confocal microscope. During imaging, cells were placed in a prewarmed humid chamber heated to 37 °C with 5% CO_2_. The cells were first starved with 1 ml of serum-free McCoy’s 5A medium and then treated with 1 ml of 007-AM (5μM) containing McCoy’s 5A medium. NIS-Elements Software was used to set time-lapse capture of a single field of view every 10 seconds for 10 min before and after the addition of 007-AM.

### Examining EPAC1 and RanBP2 interaction by immunoaffinity pulldown analyses

Approximately 2.6 ×10^6^ HEK-293 cells seeded on each 10 cm culture plate were transfected with GFP-tagged full-length wild-type EPAC1, EPAC1(SIM5A)-GFP, EPAC1(R279E)-GFP, EPAC1(1–330)-GFP, or EPAC1(331–881)-GFP for 24 hours. The cells were rinsed with PBS and lysed in ice-cold lysis buffer containing 50 mM Tris (pH 7.5), 150 mM NaCl, 1.5 mM MgCl_2_, 0.5 mM EDTA, 20 mM NEM, 1 mM phenylmethylsulfonyl fluoride (PMSF), 1% NP-40, and 1× cOmplete Protease Inhibitor Cocktail on ice for 5 to 10 min. Cell lysates were harvested by centrifugation at 16,000 g for 15 min to remove cell debris. Cell lysates with an equal amount of total cellular proteins (1 mg) were incubated with 1 μg of anti-GFP antibody (MilliporeSigma, Catalog no. 11814160001) with gentle mixing at 4 °C for 2 h. Protein A/G PLUS-Agarose beads (20 μl), equilibrated in lysis buffer, were added to the sample mixtures and incubated at 4 °C with gentle mixing for 2 h. The agarose beads were collected by centrifugation at 1000 g for 3 min and washed five times with buffer containing 50 mM Tris (pH 7.5), 150 mM NaCl, 1.5 mM MgCl_2_, 0.5 mM EDTA, 20 mM NEM, 1 mM PMSF, 0.75% NP-40, and 5% glycerol. After the final wash, the beads were resuspended in 45 μl of 2×SDS sample buffer. The GFP immunoprecipitation samples were analyzed by SDS-PAGE and immunoblotting using anti-RanBP2 and anti-EPAC1 antibodies.

To probe the effect of EPAC1 activation on RanBP2 and EPAC1 interaction, ∼ 4.6 ×10^6^ HEK293/EPAC1-V5-APEX2 cells seeded in two 10 cm dishes were starved with serum-free DMEM for 1 hour, followed by the treatment of 5 μM 007-AM or vehicle for 30 min. Cells were lysed in the same lysis buffer as described above, and cell lysates were harvested after centrifugation. Cell lysates with an equal amount of protein (800 μg) were incubated with 20 μl of V5-Trap® Agarose equilibrated in lysis buffer. After gentle mixing at 4 °C for 2 hours, V5-Agarose beads were collected with a magnet according to the manufacturer’s instructions and then washed five times with the same wash buffer as described above. After the last wash, the beads were resuspended in 45 μl of 2×SDS sample buffer. The V5 immunoprecipitation samples were analyzed by SDS-PAGE, followed by immunoblotting using anti-RanBP2 and Ran antibodies.

### Cellular Rap-GTP pulldown assay

The cellular activities of Rap1 and Rap2 in HEK293 cells ectopically expressing EPAC1-GFP or EPAC1(SIM5A)-GFP were assessed using a glutathione S-transferase fusion of the Rap1-binding domain of RalGDS as described earlier (14). Briefly, HEK293T cells at 50% confluence were transfected with pEYFP-EPAC1 or pEYFP-EPAC1(SIM5A) vector for 24 hours. Cells were starved with serum-free DMEM for 1 h followed by 007-AM (5 μM) or vehicle treatment for 30 min. After two washes in PBS, the cells were lysed in a buffer containing 50 mM Tris (pH 7.5), 150 mM NaCl, 2.5 mM MgCl_2_, 0.5% Na-deoxycholate, 0.1% SDS, 20 mM NEM, 1 mM PMSF, 1% NP-40, and 1× EDTA-free protease inhibitors. Cell lysates with equal total protein were mixed with 30 μl of glutathione-Sepharose beads with 30 μg of glutathione S-transferase-RalGDS-Rap1-binding domain bound and incubated at 4 °C for 2 h with gentle agitation. Following five washes in lysis buffer, the beads were suspended in 45 μl of 2×SDS sample buffer. 10–15 μl of protein samples were loaded onto a 15% SDS-PAGE and analyzed with Western blot using Rap1 and Rap2 specific antibodies.

### Cellular SUMOylation analysis

Cellular SUMOylation was monitored as previously described (38). Briefly, HEK293/EPAC1-V5-APEX2 or HEK293/EPAC1(SIM5A)-V5-APEX2 seeded in 12-well plates at 90% confluence were treated with EPAC-specific agonist 5 μM 007-AM or heat shock at 43°C for 30 minutes. After washing with PBS, the cells were lysed in 2×SDS sample buffer [50 mM tris (pH 6.8), 2% SDS, 0.1% bromophenol blue, 3% 2-ME, and 10% glycerol] with protease inhibitors and 20 mM NEM. Total cell lysates were collected and sonicated on ice using 15-W power output for three to four cycles of 5 seconds, with 5-s rests in between until completely soluble. After heat denaturation at 95°C for 5 min, the samples were subjected to SDS-PAGA and immunoblotting analysis with a monoclonal antibody SUMO2/3 antibody (1:3000 dilution).

### Immunoblotting analysis

Samples with equal total proteins in SDS sample buffer and boiled at 95 °C for 5 minutes were first separated by SDS-PAGE gel electrophoresis. After electrophoresis, gel images were captured using the ChemiDoc Touch Imaging System (Bio-Rad) to quantify total protein loading prior to transfer onto PVDF membranes. The membranes were then blocked with 5% nonfat milk in TBST for 1 hour, followed by overnight incubation at 4 °C with primary antibodies diluted in the same blocking buffer. Primary antibodies used in this study are Anti-EPAC1 antibody 5D3 (Cell Signaling Technology, Catalog no. 4155, 1:3000 dilution), Anti-RAN polyclonal antibody (Proteintech Group, Inc, Catalog no. 10469-1-AP, 1:3000 dilution), Anti-RanBP2 antibody (Santa Cruz Biotechnology, Inc., Catalog no. sc-74518, 1:1000 dilution), Anti-RanGAP1 antibody (Cell Signaling Technology, Catalog no. 36067, 1:3000 dilution), Anti-Rap1 antibody (Santa Cruz Biotechnology, Inc., Catalog no. sc-65, 1:1000 dilution), Anti-Rap2 antibody (BD Biosciences, Catalog no. 610215, 1:2000 dilution), and Anti-SUMO2/3 antibody (MBL Life Science, Catalog no. M114-3, 1:3000 dilution). The blots were then incubated with horseradish peroxidase-conjugated secondary antibodies (Bio-Rad) and detected using Amersham ECL Prime Western Blotting Detection Reagent (GE Healthcare Life Sciences, catalog no. 45-002-401). Chemiluminescence signals were captured with the ChemiDoc Touch Imaging System (Bio-Rad) and quantitated using ImageJ software. The signal from each specific blot was first normalized to the corresponding total protein loading. The final immunoblotting results were then expressed as the ratio of the normalized treatment signal to the normalized control signal.

### In vitro guanine nucleotide exchange factor (GEF) activity assay of EPAC proteins

In vitro EPAC GEF activity was measured as previously described using purified Rap1B(1–167) loaded with BODIPY-GDP (40). The assay was performed using 500 nM Rap1b-BODIPY-GDP and 200 nM EPAC1 or EPAC2 in buffer containing 50 mM Tris-HCl pH 7.5, 50 mM NaCl, 5 mM MgCl_2_, and 50 μM GDP, in the presence or absence of RanBP2/ZFD and at the indicated concentrations of cAMP at room temperature using 96-well plates (Corning Costar 3915). The exchange reaction was monitored using a FlexStation 3 Plate Reader (Molecular Devices) with the excitation and emission wavelengths set at 485 and 515 nm, respectively.

### Statistical Analyses

Results are presented as mean ± S.D. with N as number of independent biological replicates. A Student T-test was used to compare two groups of equal variances. One-way ANOVA with a Bonferroni post hoc test was used to compare groups of three or more with normal distributions. A p-value of less than 0.05 was considered statistically significant.

## Acknowledgments

We would like to thank Drs. Mary Dasso and Maia Ouspenskaia at the National Institute of Child Health and Human Development for providing AID::Nup358 HCT116 cells. This work was supported by grants from the National Institutes of Health R35GM122536, R01EY033319, and R01DK136462. The funders had no role in the study design, data collection and analysis, publication decision, or manuscript preparation.

## Author Contributions

Wenli Yang: Methodology; Investigation; Validation; Formal analysis; Data Curation; Writing-Method, Review & Editing; Visualization. Fang Mei: Methodology; Investigation; Validation; Formal analysis; Data Curation; Writing-Review & Editing; Visualization. Wei Lin: Methodology; Investigation; Validation; Data Curation; Writing-Method. Jason Lee: Methodology; Investigation; Validation; Writing-Method, Review & Editing. Si Nie and Chris Biley: Resources-key reagents. André Hoelz: Resources-key reagents; Writing-Review & Editing. Xiaodong Cheng: Conceptualization; Methodology; Investigation; Validation; Formal analysis; Resources; Data Curation; Writing-Original draft preparation, Review & Editing; Visualization; Supervision; Project administration; Funding acquisition.

## Competing Interest Statement

The authors declare no competing financial interests.

## Data Availability

All data are contained within the manuscript and Supporting Information.

## Notes

### Competing Interest Statement

The authors have declared no competing interest.

