## Supplemental Figures for "A SUMO-interacting motif in the guanine nucleotide exchange factor EPAC1 is required for subcellular targeting and function"

* Xiaodong Cheng

**Keywords:** cAMP; SUMOylation; nuclear pore complex; nuclear envelope; RanBP2/Nup358.

### Supplementary Figures

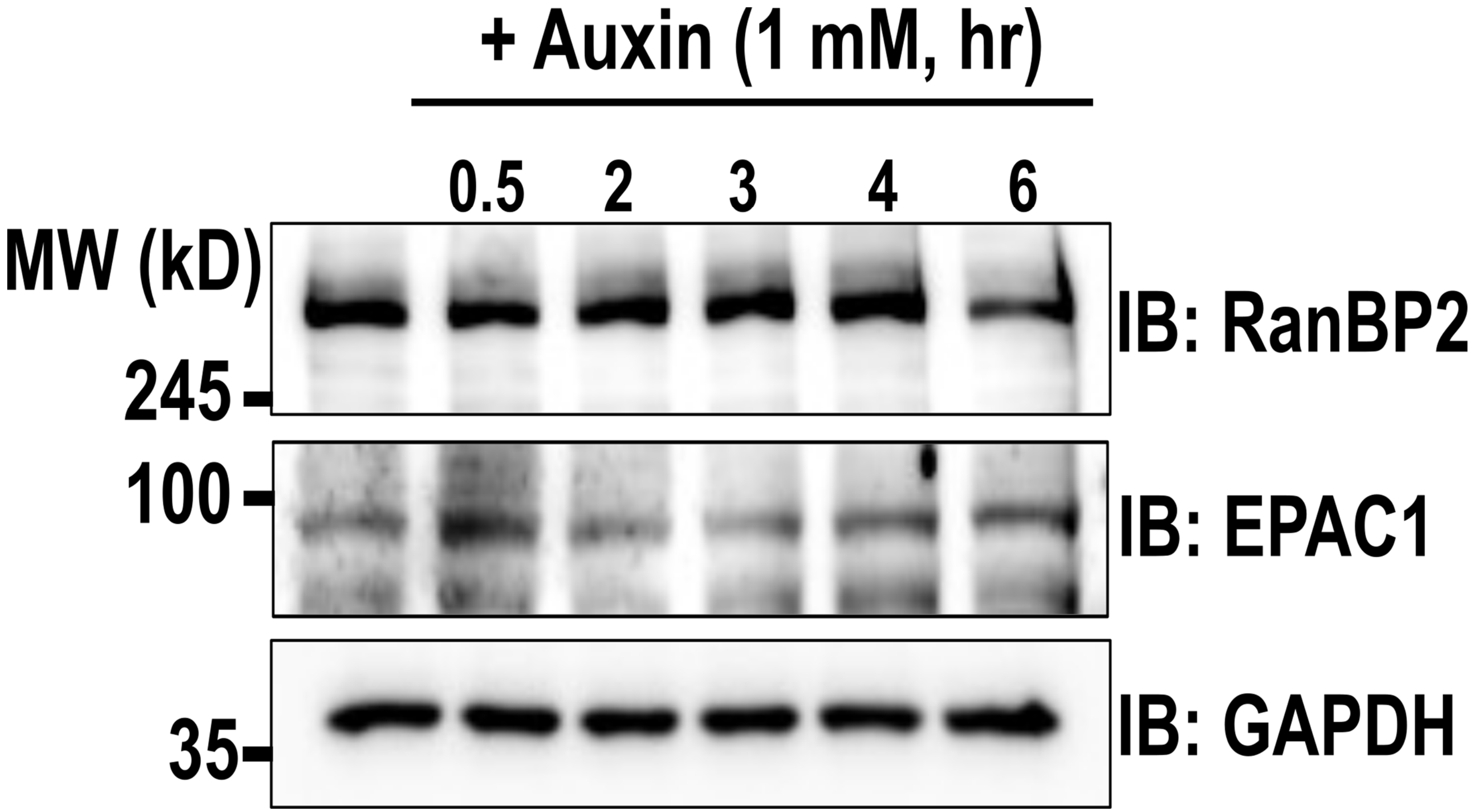


**Figure S1. Effects of Auxin treatment on endogenous cellular RanBP2 and EPAC1 in HCT116 cells.** The cellular levels of RanBP2, EPAC1, and GAPDH probed by immunoblotting analysis in HCT116 cells treated with 1 mM auxin at 37 °C for various times.

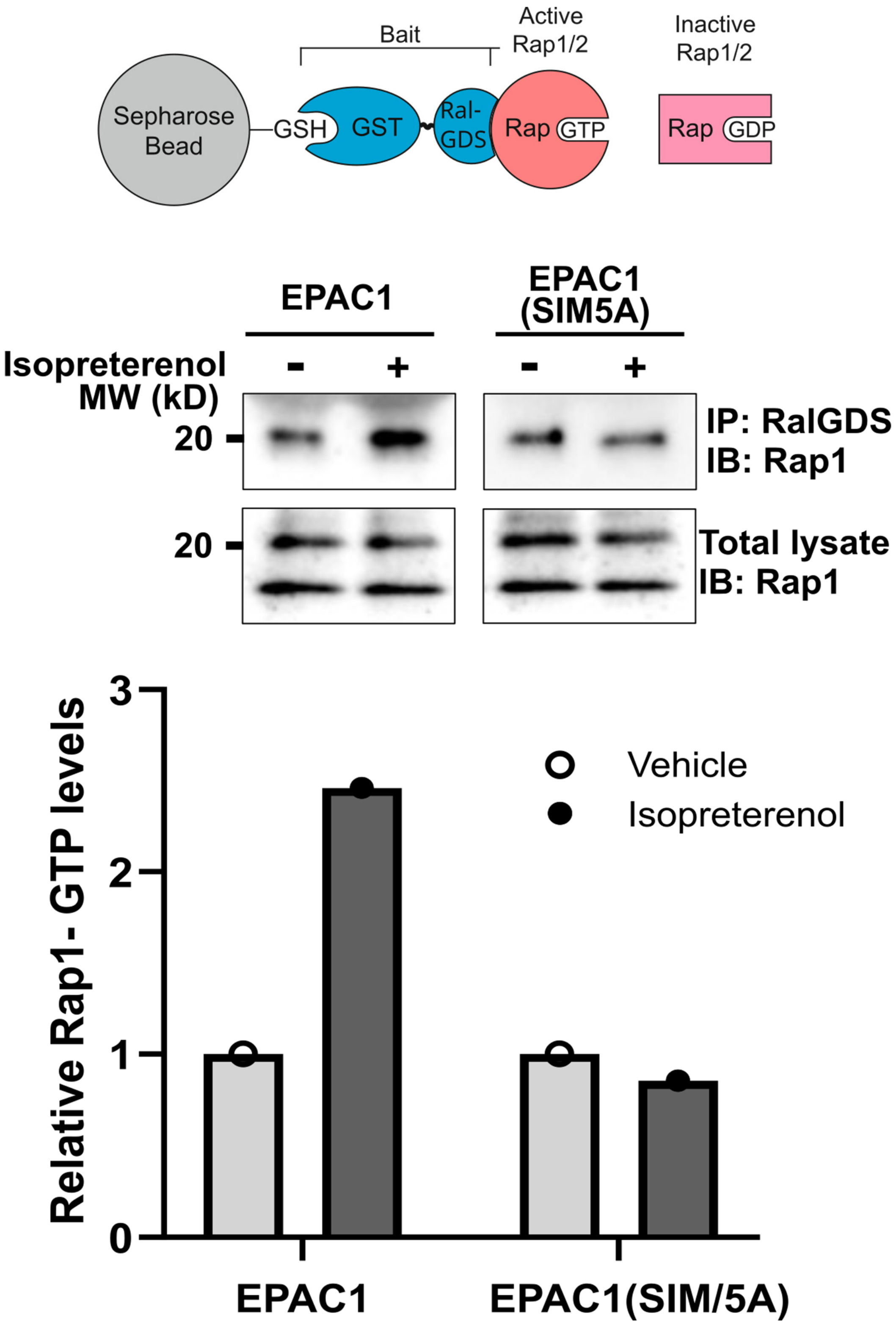


**Figure S2. EPAC1 SIM mutation interferes with cAMP-induced cellular activation of Rap small GTPases.** Levels of cellular Rap1-GTP in HEK293 cells ectopically expressing EPAC1-EYFP or EPAC1(SIM5A)-EYFP in response to 20 µM isoproterenol treatment for 30 min.

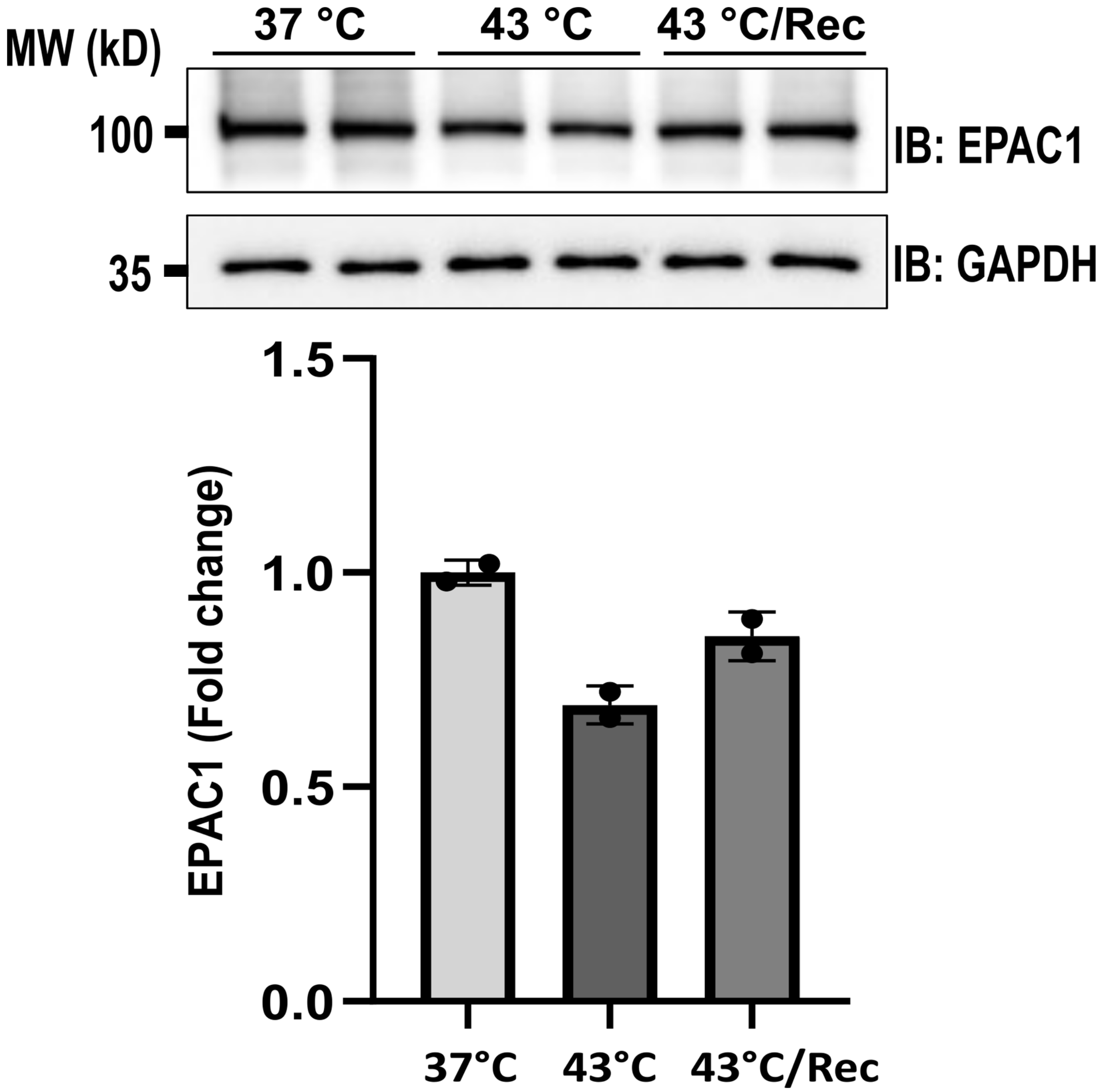


**Figure S3. Effect of heat shock on cellular EPAC1.** Levels of cellular EPAC1-APEX2 in HEK293 cells ectopically expressing EPAC1- APEX2 in response to heat shock (43 °C, 30 min) and heat shock plus recovery (37 °C, 2h).
